# A one-step biofunctionalization strategy of electrospun scaffolds enables spatially selective presentation of biological cues

**DOI:** 10.1101/850875

**Authors:** Paul Wieringa, Andre Girao, Roman Truckenmuller, Alexander Welle, Silvestro Micera, Richard van Wezel, Lorenzo Moroni

## Abstract

To recapitulate the heterogeneous complexity of tissues in our body with synthetic mimics of the extracellular matrix (ECM), it is important to develop methods that can easily allow the selective functionalization of defined spatial domains. Here, we introduce a facile method to functionalize microfibrillar meshes with different reactive groups able to bind biological moieties in a one-step reaction. The resulting scaffolds proved to selectively support a differential neurite growth after being seeded with dorsal root ganglia. Considering the general principles behind the method developed, this is a promising strategy to realize enhanced biomimicry of native ECM for different regenerative medicine applications.

## Introduction

The intrinsic heterogeneity of the body poses a great challenge when attempting to restore or replace lost functions. An emerging perspective in both biological sciences and tissue scaffold design is the influence of the microenvironment on cellular and tissue function^1^. While the traditional 2D culturing environment fails to replicate the complex and multifaceted 3D environment of the extracellular matrix (ECM), many traditional biofabrication strategies also do not capture the nuanced heterogeneous complexity of native cellular environments in a controlled manner. This limits the possibility of inducing relevant cell responses, preventing the adequate study of more complex biological systems and posing a barrier to further tissue scaffold development. In response, 3D culturing environments have been developed that exhibit controlled patterns of biologically relevant cues, including the immobilization of cell adhesion molecules in heterogeneous 3D patterns^2^ and the formation of complex 3D structures^3,4^.

Of the various scaffold fabrication strategies available, electrospinning (ESP) is often cited as a promising method of recreating the fibrous structure of the natural extracellular matrix (ECM)^5^. ESP scaffolds have proven to be particularly promising for neural tissue engineering applications^6^. By changing various parameters, this process can form fibers from tens of nanometers to tens of micrometer and can produce both randomly or oriented patterns of fibers. However, these fibers must often be modified to present cell adhesion moieties or other cues in order to better approximate the native ECM^7^. These are typically affixed to the fiber surface by the physical adsorption of proteins, chemical conjugation of biomolecules, or by incorporating ECM proteins, such as collagen, into to the polymer solution. While physical adsorption is relatively unstable, a blended fiber can require a large amount of protein mixed into the organic solvent of the polymer solution. This mixing potentially denatures the protein and renders it non-functional; as a result, this approach is expensive and potentially ineffective. The bioconjugation approach typically requires the surface of ESP fibers to be modified with reactive groups; fibers are then exposed to a solution of biomolecules, which chemically bind to the reactive groups they encounter. This method remains an efficient and effective means of fiber functionalization. However, this approach homogenously functionalizes the entire scaffold structure and cannot be used to impart a spatially defined pattern onto the fibrous construct.

Spatially selective patterns can also be achieved by photolithographic means, though this can lead to inefficient conjugation and polymer degradation^8–11^. Furthermore, conjugation only occurs on surfaces that are efficiently exposed to radiation, limiting this application to only the superficial ‘visible’ regions of 3D fibrous scaffolds. Here, we present an approach to create ‘pre-functionalized’ ESP fibers, whereby poly(ethylene glycol) (PEG) chains homofunctionalized with reactive groups are introduced to the polymer solution of a chosen biomaterial *a priori* the ESP process (Figure 1). Three different reactive groups are characterized: (i) non-selective succinimidyl valerate (SVA, an *N*-Hydroxysuccinimide variant) for amine conjugation; (ii) a more selective Thiol-Maleimide conjugation approach; and (iii) a bio-orthogonal Alkyne-Azide ‘click chemistry’ strategy. The use of small chain PEG as a carrier results in the presentation of functional groups on the fiber surface without additional processing required, representing a simple method for achieving functionalized fibers.

**Figure 1.**
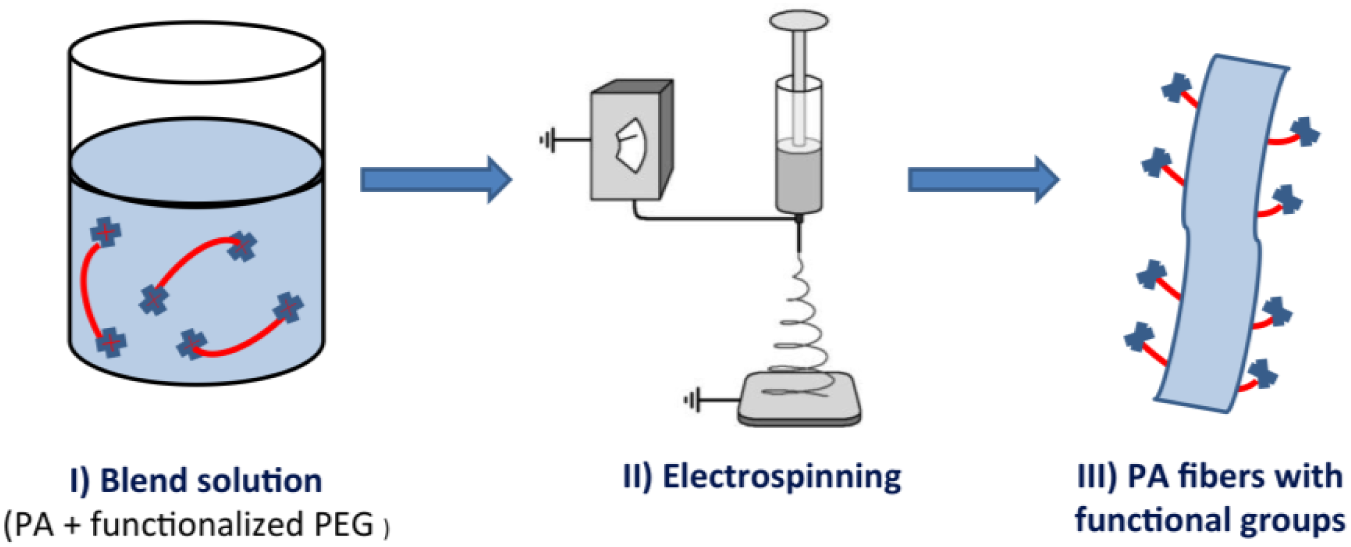
Preparation of the functionalized ESP fibers. The blended ESP polymer solution is first prepared (I), with the addition of short polymer chains (red), homofunctionalized with reactive species (blue crosses), to the bulk polymer solution. This solution is then used in the electrospinning process (II), where the resulting fibers exhibit reactive groups on the fiber surface (III).

When the different fibers are fabricated into a single construct, this provides a powerful method to selectively functionalize specific subsets of fibers within a scaffold. Combined with patterned ESP techniques, this creates a spatially defined pattern of functionalized fibers that ensures the entire fiber surface is activated. We verified the ability to create spatially defined functionalization within a 3D fibrous architecture and examine neurite growth on scaffolds functionalized with the peptide sequence H-Gly-Arg-Gly-Asp-Ser-OH (GRGDS) or the laminin-derived sequence H-Arg-Asn-lle-Ala-Glu-lle-lle-Lys-Asp-lle-OH (RNIAEIIKDI, p20)^12^. Homogeneously functionalized scaffolds successfully support neurite growth, with no difference in growth observed between peptides. However, a scaffold functionalized with both peptides in a spatially defined manner is shown to produce spatially modulated neurite growth, underscoring the value of employing complex ECM-mimicking environments to further elucidate cell behavior.

## Materials and Methods

### Polymer solutions preparations

300PEOT55PBT45, known commercially as PolyaActive™ 30055/45 (PA), a block copolymer of poly(ethylene oxide terephthalate) (PEOT), and poly(butylene terephtalate) (PBT), was provided by PolyVation B.V. (Groningen, The Netherlands). The chemical composition is represented by the notation aPEOTbPBTc, where a is the molecular weight, in g mol^−1^, of the starting PEO segments, (C_2_H_4_O)_n_, used in the polymerization process, whilst b and c are the weight ratio between PEOT and PBT blocks, respectively. Based on the manufacturer’s data n=7 for the most abundant EO oligomer length was calculated. PA was used as main polymer, prepared as a 20% w/v solution dissolved overnight in a mixture of chloroform (CHCl_3_) and 1,1,1,3,3,3-hexafluoro-2-propanol (HFiP) at a volume ratio of 7:3. The pre-functionalized agent was added 4 hours before ESP under magnetic stirring at either a 5% or 2% w/v concentration. The three different prefunctionalization agents trialed in this study were PEG-bis(SVA), PEG-bis(Alkyne) and PEG-bis(Thiol) 5000 MW, provided by Laysan Bio (Alabama, USA). The same solutions were prepared for thin film formation time-of-flight secondary ion mass spectrometry (ToF-SIMS).

### Electrospinning

Electrospinning was performed with a customized environmental chamber, maintained at a temperature of 25 °C and 30% humidity. The polymer solution was loaded into a syringe, mounted into a syringe pump and connected to a spinneret needle via Teflon tubing to supply solution at a flow rate of 1 ml/hr. The spinneret was mounted to a parallel plate, all of which was held at 20 kV and positioned 20 cm above the grounded collector. The collector was a gap electrode arrangement, with 2 mm wide electrodes and a 1 cm gap, producing aligned fibers across the gap. ESP fibers were collected onto mesh support rings, 15 mm outer diameter and 12 mm inner diameter, placed across the gap. ESP was performed for 1 minute to achieve a sufficient fiber density. Heterogeneous scaffolds were produced via the Tandem ESP (T-ESP) technique^13^. Briefly, the same electrode setup was employed but the single spinneret was replaced with two needles supplying different solutions mounted 1 cm apart.

### ToF SIMS

Separate samples were prepared on ∼0.5 cm^2^ segments of silicon wafer coated via evaporation of fluorooctyltrichlorosilane (FOTS, Sigma Aldrich) in an enclosed chamber. Thin film control samples of polymer were prepared by spin coating (2000 rpm, 1 minute) the polymer solution on to wafer segments. Silicon wafer segments were also prepared with ESP fibers by placing them between the gap of the target electrode. The same spinning parameters were employed as before.

*ToF-SIMS* was performed on a TOF.SIMS5 instrument (ION-TOF GmbH, Münster, Germany) at the Institute of Functional Interfaces, Karlsruhe Institute of Technology (KIT). The spectrometer was equipped with a Bi cluster primary ion source and a reflectron type time-of-flight analyzer. UHV base pressure was < 5×10^−9^ mbar. For high mass resolution the Bi source was operated in the “high current bunched” mode providing short Bi_1_^+^ or Bi_3_^+^ primary ion pulses at 25 keV energy and a lateral resolution of approximately 4 μm. The short pulse length of 1.1 to 1.3 ns allowed for high mass resolution. The primary ion beam was rastered across a 500×500 μm^2^ field of view on the sample, and 128×128 data points were recorded. Primary ion doses were kept below 10^11^ ions/cm^2^ (static SIMS limit). If charge compensation was necessary an electron flood gun providing electrons of 21 eV was applied and the secondary ion reflectron tuned accordingly. Spectra were calibrated on the omnipresent C^-^, C_2_^-^, C_3_^-^, or on the C^+^, CH^+^, CH_2_^+^, and CH_3_^+^ peaks. Based on these datasets the chemical assignments for characteristic fragments were determined.

For high lateral resolution imaging the primary ion source was operated in non-bunched fast imaging mode. Here, only nominal mass resolution was obtained but the lateral resolution of the instrument is in the range of 150 nm. Therefore, peaks like S^-^ and HS^-^ were used for imaging since both peaks do not show other signals at the same nominal mass (m/z).

### Fluorescent Label Conjugation

Reactivity and availability of functional groups were confirmed by the conjugation of fluorescent dyes with complimentary binding motifs. SVA was conjugated to FITC-labeled BSA (Sigma) at 10 μg/ml in a PBS solution at pH 7.2. The thiol functional group was conjugated with a Dylight 488 maleimide probe (Invitrogen), which was applied at a concentration of 10 μg/ml in PBS. The click chemistry Alkyne functional group was conjugated with azido-labeled Megastokes 673 dye (Sigma) at a concentration of 10 μg/ml in PBS with 25 mM ascorbic acid and 2.5 mM copper (II) sulfate. Dye solutions were applied overnight. A T-ESP scaffold was simultaneously conjugated with a solution containing both the maleimide and azide probes (same concentrations). Substrates were washed 3 times with a tris buffered solution (TBS), pH 8, followed by a 1 hour wash with a TRIS-based EDTA solution, pH 9, and then demineralized water (dH_2_O) three times. To facilitate spatial analysis of observed neurite growth, T-ESP scaffolds conjugated the azide-Megastokes dye to the alkyne fibers as described above, but with a 50 μg/ml dye concentration incubated for 2 days.

### Peptide preparation

Both the click chemistry and the thiol strategies require functionalization of the biomolecule of interest with the complimentary binding partner. Peptide solutions of H-Gly-Arg-Gly-Asp-Ser-OH (GRGDS) and H-Arg-Asn-Ile-Ala-Glu-Ile-Ile-Lys-Asp-Ile-OH (RNIAEIIKDI; p20), (Bachem) were prepared at 5 mg/ml in PBS. These were activated with a maleimide and an azide, respectively, via amine-reactive linker molecules of NHS-peg4-Azide (Jena Bioscience) and sulfosuccinimidyl-4-(*N*-maleimidomethyl)-cyclohexane-1-carboxylate (Sulfo-SMCC, Thermo Scientific). Maleimide and azide linkers molecules were first dissolved in DMSO and then added to peptide solutions for a final linker concentration of 20 mM and 50 mM, respectively, representing a 5 times molar excess with respect to peptide concentration. Solutions were maintained at room temperature for 4 hours. The SVA conjugating strategies did not require any pre-preparation of the proteins to be applied.

### Scaffold Preparation

Glass cover slips (14 mm in diameter) were placed in a 24 well plate, followed by the ESP fibers collected on mesh rings; the glass cover slip provided a necessary mechanical support for subsequent removal and imaging of the electrospun mesh. A Viton rubber ring (Eriks B.V., The Netherlands) was inserted into the well to hold the scaffold in place. These constructs were placed inverted with respect to their fabrication orientation, such that the first deposited fibers were facing upwards; the mesh ring provides a physical barrier between the ESP fibers and the Viton ring, improving handling during subsequent processing of cell culture samples. For homogenously functionalized scaffolds, 150 μl of peptide solution was applied at a concentration of 0.5 mg/ml in PBS. Heterogeneous scaffolds were prepared using mixed peptide solutions at a concentration of 0.5 mg/ml of each peptide. Solutions involving click chemistry also had an additional 25 mM of L-ascorbic acid and a 2.5 mM Cu_2_SO_4_ (Sigma Aldrich). Solutions were left over night to allow for conjugation. Similar to the fluorescent label conjugation, substrates were washed 3 times with a tris buffered solution (TBS), pH 8, followed by a 1 hour wash with a TRIS-based ethylenediaminetetraacetic acid (EDTA) buffer (Klinipath B.V., The Netherlands), pH 9, and then dH_2_O three times.

### Cell Culture

Scaffolds were sterilized with 70% ethanol for 30 minutes, which was later left to evaporate. Scaffolds were washed 3 times with sterile PBS, followed by a one-time wash in Neuralbasal® culture medium (Invitrogen) supplemented with B27 supplement, 0.5 mM l-glutamine, 10 U/ml of penicillin/streptomycin (all from Invitrogen) and 10 ng/ml of NGF (Sigma Aldrich). Culture plates were left to warm with 150 μl of culture medium per well in an incubator at 37 °C and 5% CO_2_. Dorsal root ganglia (DRGs) were extracted from 2-day-old post-natal Sprague-Dawley rat pups. All procedures followed national and European laws and guidelines and were approved by the local ethical committee. Briefly, rats were sacrificed by cervical dislocation under general anaesthesia (4% Isoflurane) and then decapitated. Individual ganglia were removed from the spinal column and nerve roots were stripped under aseptic conditions with the aid of a stereomicroscope. Connective tissue was removed and the DRGs were cut to expose the enclosed cells. One DRG was placed in the center of each ESP fiber scaffold; particular care was taken to ensure the DRG was placed in the overlapping region of the T-ESP scaffolds. Cultures were maintained for 5 days, with medium refreshed every other day.

### Immunohistochemistry

Cells were first cooled to 4 °C, followed by the addition of an equal volume of ice cold 4% w/v paraformaldehyde (PFA) in PBS, added to each well for a final concentration of 2% PFA. Fixation proceeded for 20 minutes, after which samples were washed with Tris Buffered Solution (TBS). Samples were permeabilized with 0.1% Triton-X for 15 minutes, washed 2 times with TBS and blocked with 5% normal goat serum for 1 hour. This was followed by a 16 hour incubation at 4 °C in a 1:1000 dilution of BIII-tubulin (Sigma) raised in mouse, and 1:500 S100 (Sigma) raised in rabbit in TBS with 1% normal goat serum. Samples were then washed 3 times in TBS with 1% normal goat serum, secondary antibody of anti-mouse Alexa 488 or Alexa 546 raised in goat (Invitrogen) and anti-rabbit Atto 647N raised in goat (Sigma) were applied at a dilution 1:500 for 12 hrs at 4 °C. Samples were then washed 3 times with TBS containing 1% BSA and 0.1% w/v sodium azide (Sigma Aldrich). After staining, the alkyne-containing fibers of the T-ESP scaffolds were selectively labeled as outlined in the *Fluorescent Label conjugation* section above. Samples were washed again with TBS, then removed from the well and mounted on a coverslip using 4-88 mowiol mounting fluid with 2.5% w/v 1,4-diazabicyclo[2.2.2]octane (DABCO, Sigma Aldrich) for microscopy imaging.

### Microscopy and Image Analysis

Fluorophore conjugation to fibers was imaged using a Nikon TI confocal microscope, eliminating diffractive autofluorescence. Per fluorophore, laser power was adjusted using the conjugated scaffolds as a reference. The remaining scaffolds were imaged using the same power settings to determine the degree of selective conjugation. Cell labels were imaged using a BD Pathway 435 with a 4× objective. Neurites were imaged via the Alexa 488 or Alexa 546 labels, using the ex: 482/35-em: 536/40 or ex: 543/22-em: 593/40 filter sets respectively (ex: excitation; em: emission). Schwann cells were visualized with the Atto 647N fluorophore using an ex: 628/40-em: 692/40 filter set. The use of Megastoke 678 dye made possible the visualization of alkyne fibers by using an ex: 543/22-em: 692/40 filter set. Neurite outgrowth was assessed by spatially measuring the number of pixels in radially segmented concentric bands around a DRG. Images shown were processed using the Contrast Limited Adaptive Histogram Equalization (CLAHE) filter within the ImageJ/Fiji software package, with stitched mosaic created with the Grid Stitch plug-in ^14^.

### Statistical Analysis

All statistics were performed with R statistical software (http://www.R-project.org/) and graphs were created using the Deducer plugin (by Ian Fellows, http://www.jstatsoft.org/v49/i08/). Comparisons of fiber diameters and fiber alignments were performed with a one way ANOVA followed by a *post hoc* Holm method, with significance level of p < 0.01 and a minimum of 100 fibers measured per fiber type.

## Results

Initial ESP trials incorporated functionalized PEG additives at a concentration of 5% w/v. However, the quality of the fiber deposition across the gap electrode was inconsistent. An apparent phase separation of the blended polymer solution was also observed over a 12 hour period, suggesting poor solubility of the PEG additive.

Reducing the additive concentration to 2% produced a stable polymer solution and an improved ESP fiber deposition (Figure 2). ALK and SH fibers produced fibers of 0.88±0.23 μm and 0.71±0.19 μm in diameter, with a statistically significant difference between the two populations. The SVA fibers were much larger in comparison, with a diameter of 1.56±0.47 μm. Image analysis of fiber alignments provides a coherence metric between 0 (random) and 1 (aligned). ALK and SH fibers were found to have statistically similar degrees of alignment (0.42±0.06 and 0.35±0.07, respectively), while the SVA fibers exhibited a higher degree of alignment (0.59±0.08).

**Figure 2.**
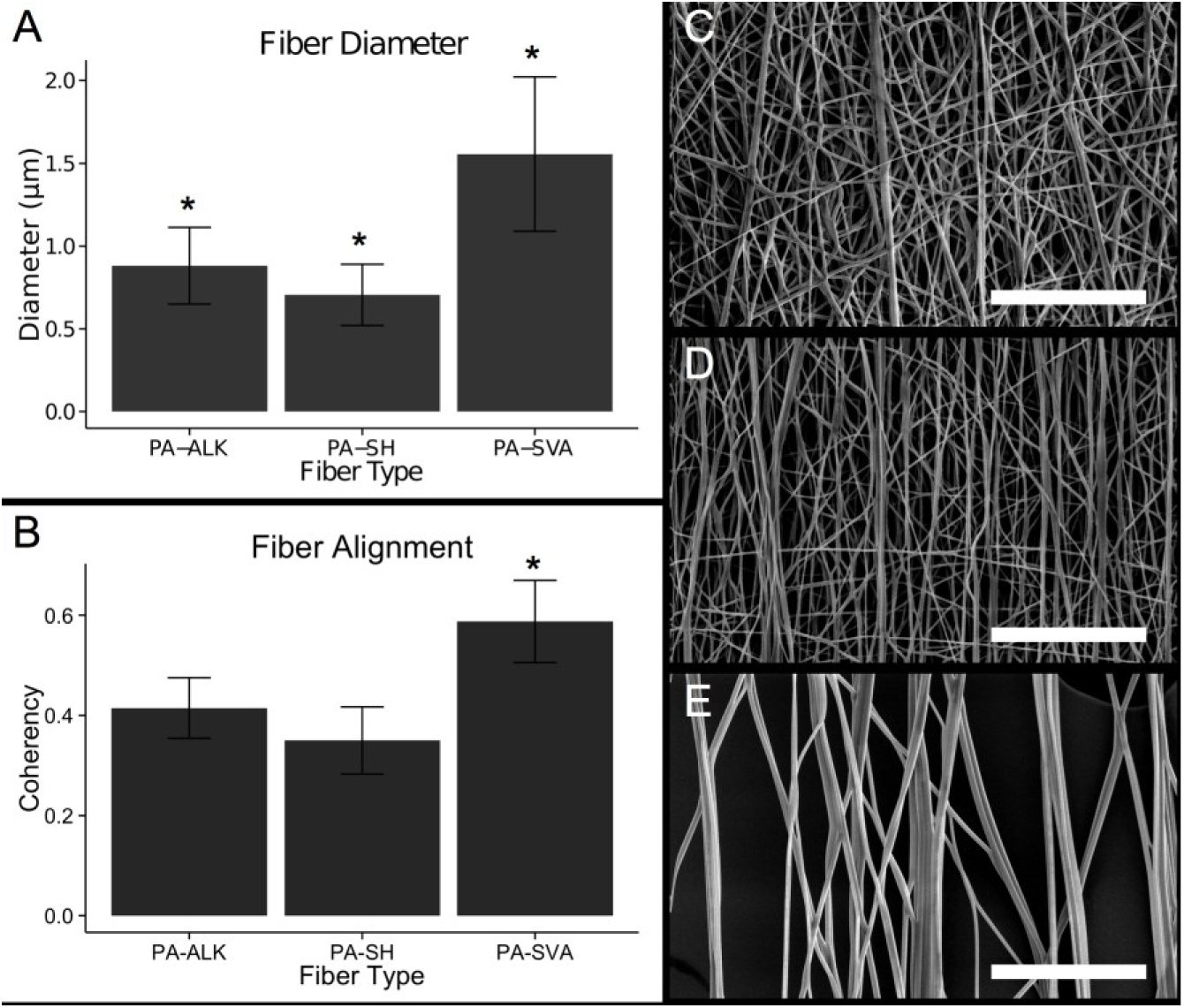
ESP fiber size and alignment. Diameters are shown for PA fibers with 2% of ALK, SH and SVA conjugation agents (A). All three diameters were found to be statistically different from each other, according to a one-way ANOVA with a post-hoc Holm-corrected pairwise comparison (p< 0.01; n=100, fibers per sample). The degree of fiber alignment was also determined (B) using coherence as a metric, where 0 is random and 1 is perfectly aligned. Similar analysis found that fiber alignments of the ALK and SH fibers were similar, while the SVA fibers were significantly more aligned (p<0.01, n =6, images per sample). Scalebars: 50 μm.

ToF-SIMS analysis provides a method to identify elemental composition of the first few nanometer of a surface, providing an effective means of assessing functional group availability on the pre-functionalized ESP fibers. Plain PA fibers were characterized by several molecular fragments based on the EO repeating units. Moreover, a rather large fragment of the PEOT block, assigned to [(C_2_H_4_O)_7_ C_7_H_5_O_2_]^-^, was detectable in negative polarity SIMS. Based on the matching ^13^C isotope pattern in the mass spectrum the multiplet would correspond to the most abundant PEOT block after cleavage of the terminal CO group and ionization by hydrogen uptake. Despite their low abundance both the 2% SVA and 2% SH fibers clearly exhibited available functional groups, being absent in plain PA samples. Strong and characteristic signals were, among others, the C_4_H_4_NO_2_^-^ fragment found for the succinimidyl group of SVA; and S^-^ together with SH^-^ in case of the thiol presenting fibers, SH. In secondary ion mass spectra recorded with high mass resolution no interfering signals were present in case of the marker signals given above. Hence, it was possible to record SIMS images with high lateral resolution in a non-bunched mode providing only nominal mass resolution as shown in Figure 3. Unfortunately, ALK fibers were not possible to be assessed because the chemical signature of the incorporated alkyne groups was not sufficiently distinct to be discerned from the bulk PA polymer.

**Figure 3.**
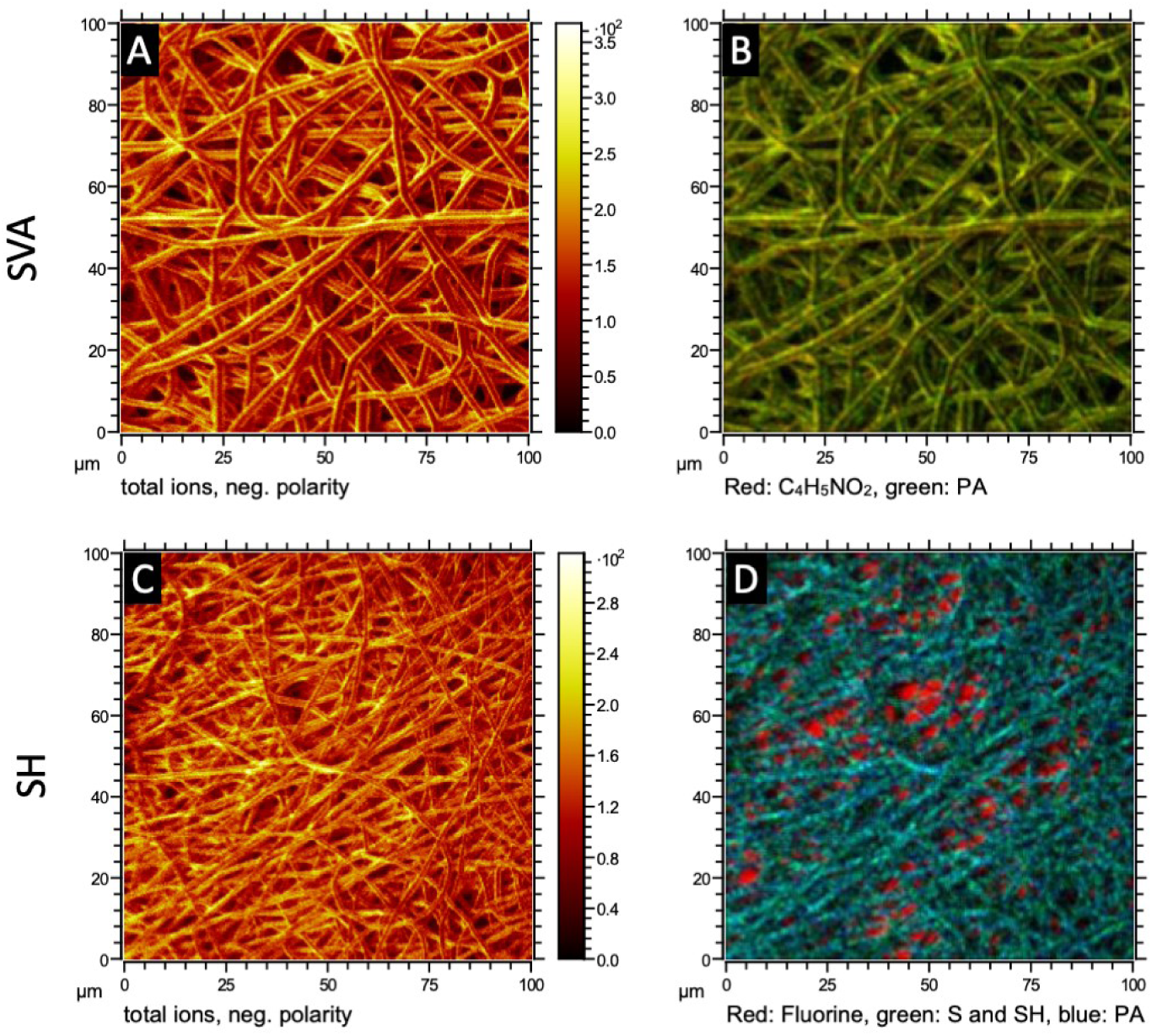
High lateral resolution ToF-SIMS imaging. (A) and (C): total ion images of SVA and SH fibers, respectively. (B): Overlay for SVA sample. Red: succinimidyl moiety C_4_H_4_NO_2_^-^, green: multiplet at 429 m/z indicating the PEOT block in PA polymer. (D): Overlay for SH sample: Red: F^-^ from the substrate, green: sum of S^-^ and SH^-^, blue: multiplet at 429 m/z indicating PEOT repeating unit in PA polymer.

To ascertain whether chemical groups on the fiber surface were available for conjugation, solutions of fluorescent probes with complementary conjugation molecules were employed; these included BSA-FITC for SVA conjugation, Dylight-488 with maleimide for SH conjugation and Megastokes 678 with an azide for alkyne (ALK) conjugation. Available groups on the fiber surface were reactive (Figure 4). The SVA fibers were shown to strongly retain the BSA-FITC protein (Figure 4A) compared to the ALK and SH fibers (Figure 4D, G), although a degree of non-specific binding could still be observed. A high degree of selective conjugation was noted for the smaller molecular probes used for the ALK fibers (Figure 4F), with no observable cross-reactivity with either the SVA fibers (Figure 4C) or SH fibers (Figure 4I). Similarly, the maleimide fluorescent probe showed a clear preference for the SH fibers (Figure 4H) compared to no observed reactivity with the SVA or ALK fibers (Figure 4 B, E).

**Figure 4.**
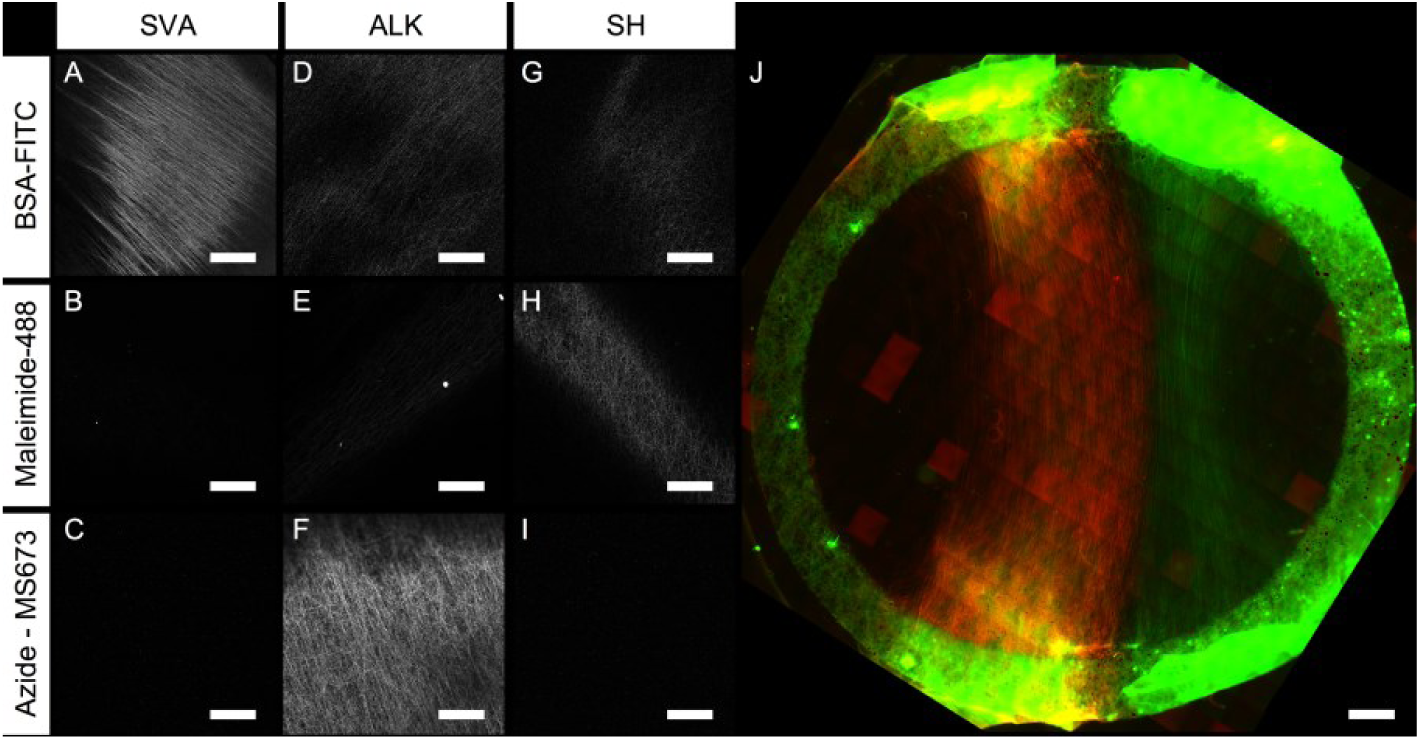
Fluorescent probe conjugation on different fiber types. The SVA additive fibers showed strong binding to BSA-FITC (A), where BSA is a representative large globular protein with available amine groups. No signal was observed when the maleimide dye (B) or azide dye (C) were applied to SVA fibers. ALK and SH fibers both experienced residual adhesion of the BSA (D, G). Both of these fibers were shown to have highly specific binding to azide-MegaStokes 673 (MS673) and Maleimide-Dylight 488 (Maleimide-488) (H, F). A T-ESP scaffold of ALK and SH fibers within a mesh frame (autofluorescent green boundary) showed selective conjugation of azide-MegaStokes 673 (red) and maleimide-Dylight 488 (green), respectively. Scale bar: A-I = 250 μm; J = 1 mm.

Tandem ESP was also employed to create a heterogeneous scaffold of two aligned fiber populations positioned next to each other with defined overlapping region. A tandem scaffold of ALK and SH fibers was exposed to a solution of maleimide-Dylight 488 and azido-MegaStokes 673, verifying that spatially selective conjugation of different fibers types is possible with such a system.

To validate these conjugation strategies, neurite outgrowth was evaluated on functionalized scaffolds. The SH and ALK fibers were selected for *in vitro* evaluation, owing to similarities of fiber diameter and alignment as well as high conjugation selectivity and similar use of activated proteins. SH fibers and ALK fibers were functionalized with the GRGDS and p20 peptides, respectively.

Neurite outgrowth on these substrates was well aligned in the direction of fiber orientation (Figure 5A, C), with no distinct differences in the length or distribution of outgrowth on fibers functionalized with either the GRGDS or p20 peptide as evidences by pixel distribution analysis of axon growth (Figure 5A, C inset). The migration of Schwann cells from the explanted tissue was also observed on both fiber substrates (Figure 5B, D). Fibers functionalized with GRGDS exhibited a higher density of Schwann cells surrounding the explanted DRG.

**Figure 5.**
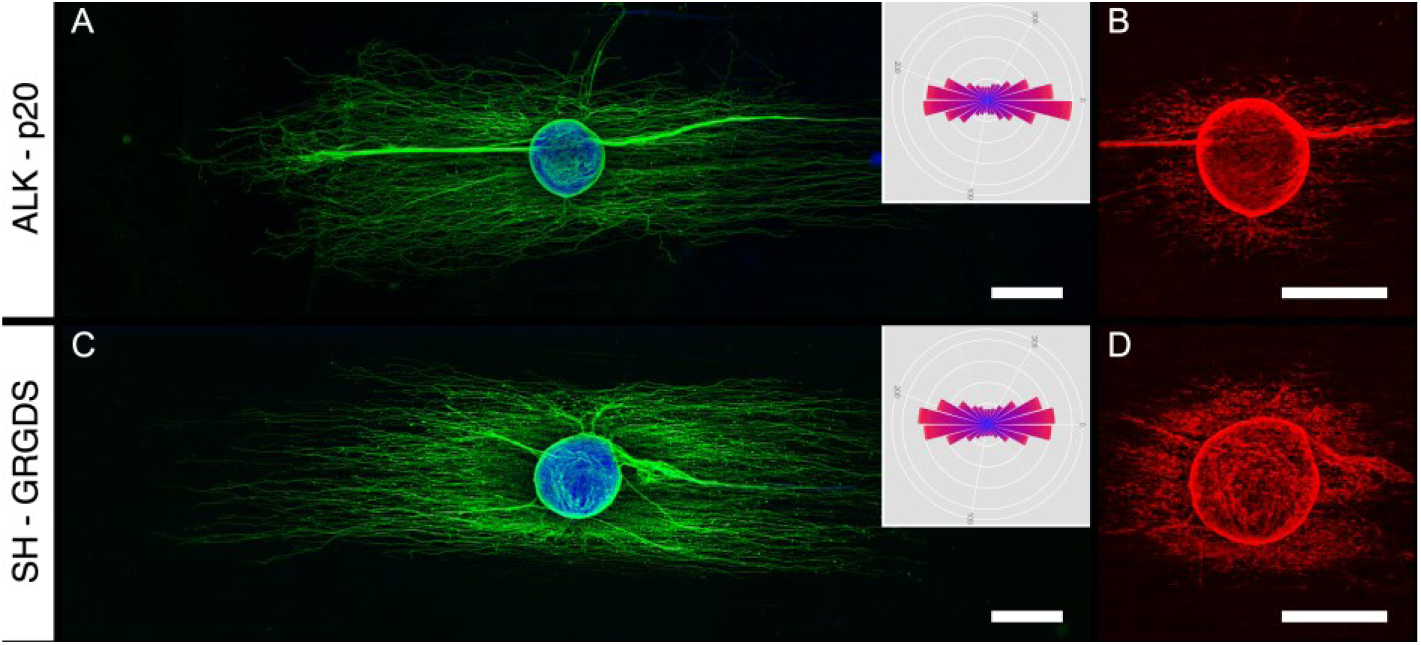
Representative images of neurite outgrowth and Schwann cell migration from explanted DRGs. Neurites outgrowth on the p20 fibers (A) or GRGDS fibers (C) was equivalent, confirmed by analyzing the neurite distribution (inset). P20- and GRGDS-functionalized fibers also stimulated Schwann cell migration from the DRG explant (B, D, respectively). Scale bars: 500 μm.

**Figure 6.**
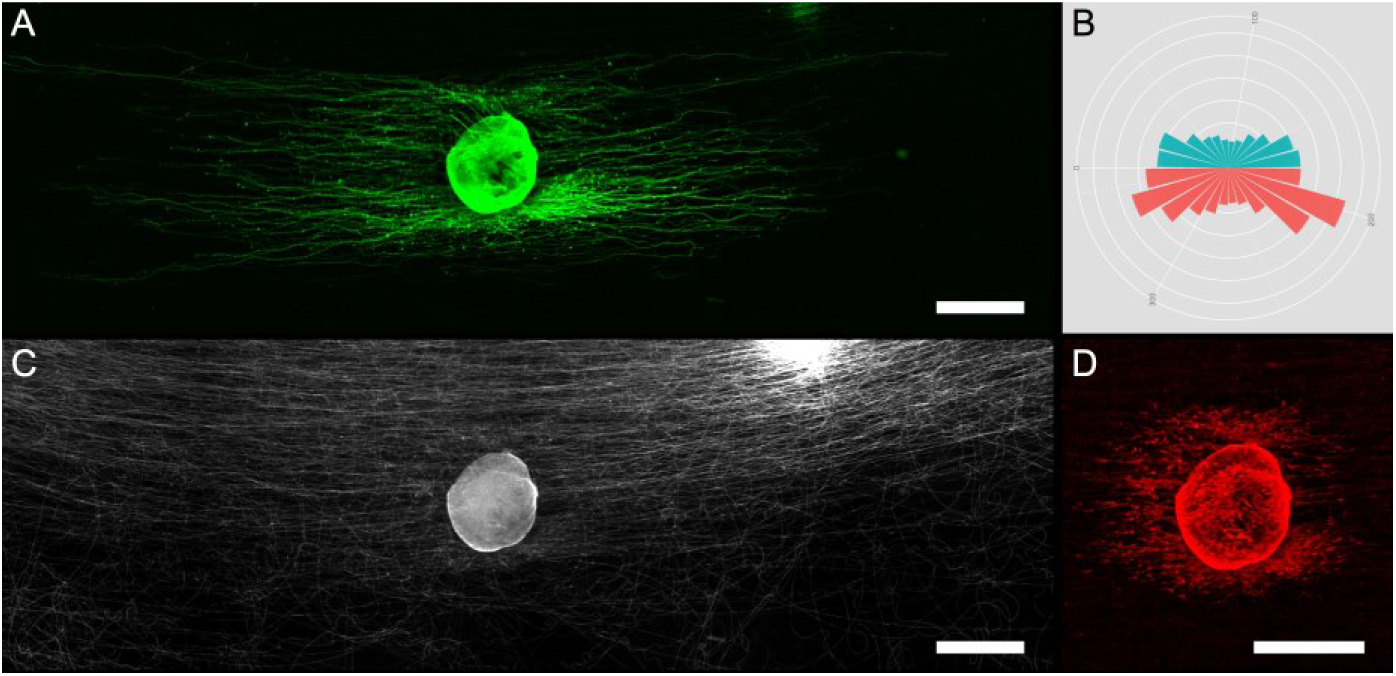
Representative outgrowth on Tandem ESP (A). The cumulative pixel intensity of neurite outgrowth (B) provided a comparison of the density of neurite growth, with the GRGDS fibers exhibiting higher density outgrowth. Fibers with p20 functionalization were visualized (C, in white) versus the region of RGD-functionalized fibers (C, black region), with the DRG placed at the nexus of the two fiber populations. Also observed was Schwann cell migration from the explanted DRG (D). Scale bars: A, C, D = 500 μm.

These peptides were also applied to tandem ESP scaffolds, creating spatially defined regions of aligned fiber with specific biomolecule functionalization. Under these conditions, a preferential growth on GRGDS-modified fibers was revealed, despite the presence of the neurite-specific p20 peptide. This is despite an apparent uniform distribution of migrating Schwann cells.

## Discussion

The goal of this study was to develop ESP fibers that could be functionalized with biomolecules without the need of post-modification by wet chemistry or photolithography. While representing a simple strategy for fiber bioconjugation, this was pursued to create fibers with orthogonal conjugation chemistries to enable the selective functionalization of specific subpopulations of fibers within a heterogeneous ESP scaffold. We were able to verify the presence of reactive species on ESP fibers and confirm the selectivity of the different chemistries employed. Applying this approach to the assembly of spatially defined heterogeneous fiber scaffolds, created via the T-ESP technique, a fibrous scaffold with spatially defined, orthogonal conjugation chemistries was created. As an example, preliminary results showed that neurite growth on such scaffolds differed compared to homogeneous scaffold and exhibited spatially modulated behavior.

The ability to conjugate biomolecules to 3D culturing platforms in a spatially defined manner represents a promising tool to study and dictate cell response. While studies of cells on 2D substrates with biomolecule patterns have produced interesting results^15,16^, planar substrates limit the applicability of these outcomes. Many of the relevant signals imparted to a cell are lost or altered when moving between a native 3D ECM and a 2D substrate. However, the creation of analogous biomolecular patterns within a 3D scaffold via spatial selectivity represents a biofabrication challenge, with emerging strategies employing two-photon activation of reactive molecules within optically-permissive hydrogel environments^2,17,18^.

Outlined in this current study is a method to spatially define bioactive moieties within a 3D fibrous scaffold, providing cells with a combination of nanotopography and a pattern of biomolecular cues. Functionalization of ESP fibers traditionally relies on wet chemistry to bestow reactive groups on the fiber surface, a method that does not permit spatially defined bioconjugation. Recently, alternative approaches to functionalize ESP fibers have been explored whereby the polymer solution was modified to include functionalized reactive groups before ESP. These approaches included conjugation chemistries similar to those employed in the current study^19–21^. However, all methods required the custom synthesis of active carrier polymer chains. In contrast, the method described in the current study provides a simple means of attaining pre-functionalized ESP fibers through the use of commercially available modified PEG chains as a carrier. ToF-SIMS analysis revealed the availability of reactive groups on the fiber surface, confirmed by the successful conjugation of fluorescent probes. This approach is compatible with many polymer preparations currently used in ESP and creates fibers nearly identical in composition while varying surface reactivity by simply modifying the used additive.

Neurite growth from an explanted rat DRG was evaluated on scaffolds conjugated with either p20 or GRGDS peptides, with no observable differences in the degree of outgrowth were observed. To showcase the utility of this approach, a heterogeneous scaffold comprised of two overlapping populations of aligned fibers was created. One fiber type incorporated the ALK additive and was subsequently functionalized with p20 using click chemistry, while the other contained the SH additive and functionalized with the GRGDS adhesion motifs via thiol-maleimide chemistry. An explanted DRG was then placed on the overlapping region between these two fiber groups. Despite the equal performance of these two peptides on homogenously conjugated scaffolds, biased neurite growth was observed on the RGD-functionalized fiber population.

RGD is a well-known cell adhesion peptide sequence, present in many ECM proteins such as fibronectin, vitronectin, collagen and laminin^22^. This adhesion molecule is known to interact with cells through a well characterized family of integrins, with clear evidence of positive influence on neurite outgrowth^23^. In contrast, p20 is a lesser known sequence found specifically on the gamma 1 chain of laminin. Identified by Liesi *et al.* as promoting neurite outgrowth^12^, it is now known to bind to the non-integrin cell membrane prion protein (PrPc). This receptor is considered a modulator of cell behavior, maintaining the destabilization of β1 integrin-related focal adhesion formation and causing localized release of Ca^2+^ ions, resulting in increased cell motility and neurogenesis^24^.

The DRG explant model used in this study also includes the complex interaction between neurons and supporting glial Schwann cells, also present in DRG explants and highly active during peripheral nerve regeneration. After injury, Schwann cells typically precede the regenerative front *in vivo*. They change phenotype to produce ECM molecules and diffusible growth factors, encouraging and guiding neurite growth^25^. Recent work in our lab has shown that Schwann cells exhibit a differential response to both the p20 and RGD peptides. While both of these peptides promoted increased metabolic and proliferative activity in comparison to non-functionalized fibers, the RGD sequence was observed to also upregulate NGF production^26^.

Within the current study, the observed bias in neurite growth is thought to be the result of a localized release of NGF by Schwann cells in response to RGD-conjugated fibers. For scaffolds that are homogeneously functionalized with either the NGF-promoting RGD or neurite promoting p20, evidence of such a localized release of NGF is lost and resulting growth is similarly homogenous. For tandem-spun scaffolds, the spatially defined distribution of different peptides likely produced a local concentration gradient. This attracted the initial outgrowth of neurites in a biased manner, with a preference for the RGD sector. This is despite virtually no difference in the distribution of Schwann cells in and around the DRG, attributed to both RGD and p20 as effective substrates for this glia subtype^26^.

When this approach is implemented within a heterogeneous scaffold, it presents a promising approach to investigate the complexity of a heterogeneous growth environment. These findings exemplify the importance of examining heterogeneity for both natural and designed ECM constructs. Though traditional homogeneous environments will continue to provide knowledge, these substrates overlook the influence of ECM components on the spatial-temporal regulation of cell behavior. Though these preliminary results are still far from achieving decisively selective cell response, the methodology proposed here provides the first step towards mimicking the complexity of the ECM heterogeneity.

## Conclusion

We introduced three facile methods for creating functionalized fibers and present the concept of combining these strategies to create an ordered, heterogeneous tissue scaffold. The use of such scaffolds reveals the synergistic influence of different peptides on the function of glia and neurite growth, revealing how such multimodal fabrication strategies might assist in both understanding the complexities of biological systems and providing tissue engineering solutions.

